# Human mismatch repair system corrects errors produced during lagging strand replication more effectively

**DOI:** 10.1101/045278

**Authors:** Maria A. Andrianova, Georgii A. Bazykin, Sergey I. Nikolaev, Vladimir B. Seplyarskiy

## Abstract

Mismatch repair (MMR) is one of the main systems maintaining fidelity of replication. Different effectiveness in correction of errors produced during replication of the leading and the lagging DNA strands was reported in yeast, but this effect is poorly studied in humans. Here, we use MMR-deficient (MSI) and MMR-proficient (MSS) cancer samples to investigate properties of the human MMR. MSI, but not MSS, cancers demonstrate unequal mutation rates between the leading and the lagging strands. The direction of strand asymmetry in MSI cancers matches that observed in cancers with mutated exonuclease domain of polymerase δ, indicating that polymerase δ contributes more mutations than its leading-strand counterpart, polymerase ε. As polymerase δ primarily synthesizes DNA during the lagging strand replication, this implies that mutations produced in wild type cells during lagging strand replication are repaired by the MMR ~3 times more effectively, compared to those produced on the leading strand.

## Results

Replication is a very accurate process. Its fidelity is achieved through three main components: base selectivity of polymerases, proofreading activity of their exonuclease domains, and repair of mismatches that escaped proofreading by mismatch repair (MMR) system ^1^. Studies in yeast indicate that the effectiveness of each of these steps depends on the mismatch type, and that MMR compensates for the infidelity of polymerases ^2,3^. According to the current model of the replication fork in a eukaryotic cell, polymerase ε preferentially copies the leading strand template, while polymerases α and δ copy the lagging strand template ^4–8^. In yeasts, different replicative polymerases possess different biases in the types of mutations they introduce, leading to differences in mismatch types between the leading and the lagging strands ^2^. These biases can be further modulated by differences in MMR effectiveness between the two strands. For example, in yeast, the 8-oxo-guanine-adenine mismatches generated during the lagging strand replication were shown to be repaired more effectively, compared to those generated during the leading strand replication ^9^. More generally, in yeast, the biases between leading and lagging strands introduced by the MMR during repair tend to be opposite to the biases introduced during replication, thus compensating for them ^10,11^.

We first asked which types of substitutions are corrected by the MMR more effectively in humans. To address this, we compared the mutational spectra in MSI and MSS cancers of two different cancer types – colon adenocarcinoma (COAD) and uterine corpus endometrial carcinoma (UCEC) (Fig. 1a, b). In line with previous studies ^12–14^, we found that MSI cancers are enriched in C to T (C→T) mutations in the Gp<u>C</u>pN context, A→G mutations in all contexts, and C→A mutations in the Cp<u>C</u>pN context (Fig. 1a, c). Therefore, these mutation types are effectively corrected by the MMR.

**Figure 1.**
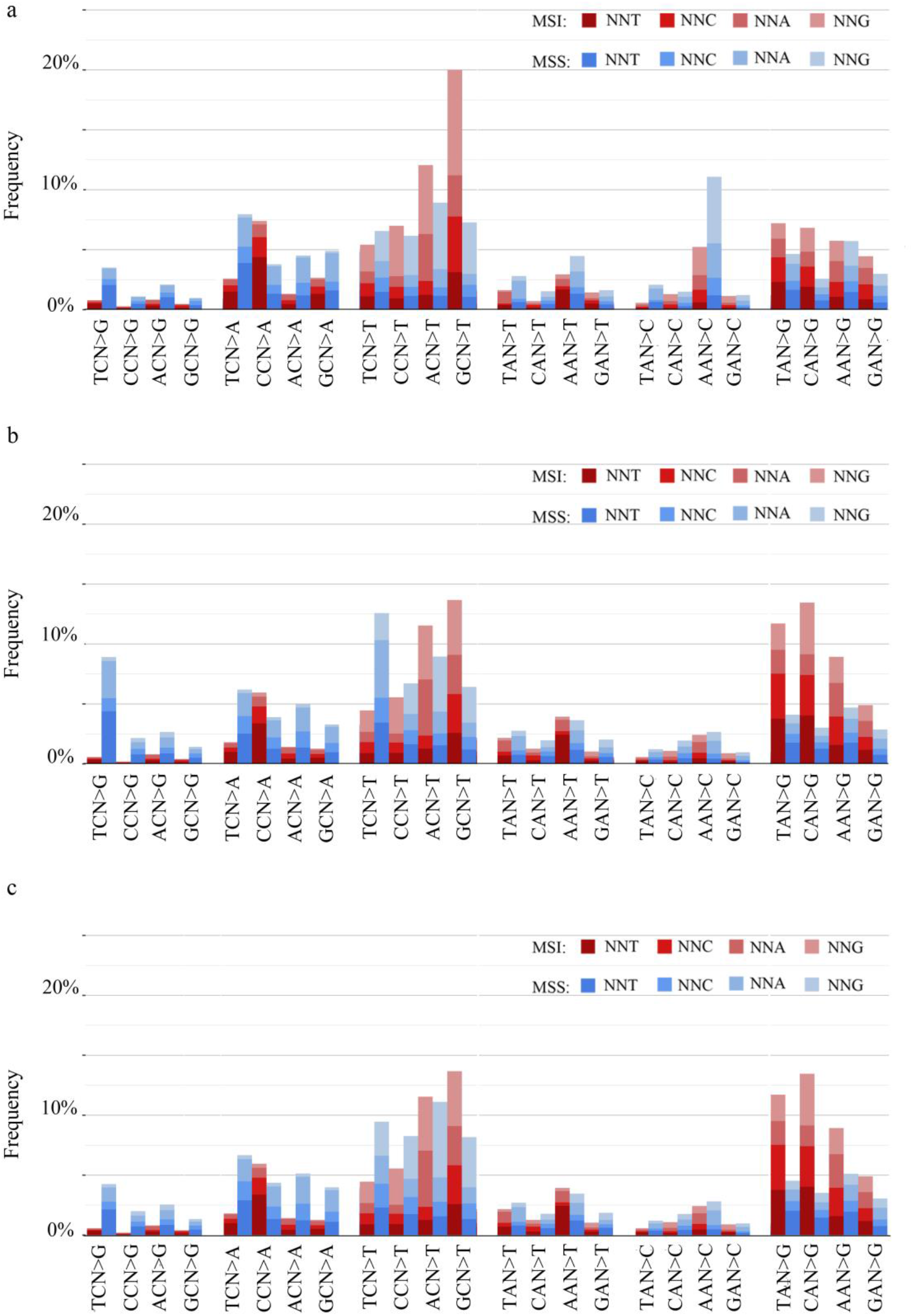
**Mutational spectrum of MMR deficient cancers**. Relative frequencies of the 96 mutation types in MSI and MSS cancers. Complementary mutation types were pooled. **a**, MSI (n = 3) and MSS (n = 8) colon adenocarcinoma samples; **b**, MSI (n = 7) and MSS (n = 19) uterine corpus endometrial carcinoma; **c**, MSI (n = 7) and MSS (n = 14) uterine corpus endometrial carcinoma excluding APOBEC enriched samples.

The mutational spectra of the two cancer types are somewhat distinct. To address these differences, we calculated the weights of mutational signatures ^13^ in the spectrum of each sample. In all MSI samples, we observed signatures of defective DNA mismatch repair (Signatures 6, 15, 20, 26). In many MSI samples of UCEC, we also observed clock-like signature 5 and other mutational signatures not related to MMR deficiency (Supplementary Table 1). Therefore, the mutational patterns in UCEC MSI cancers are slightly confounded by MMR-independent mutational processes. Among the MSS samples, a strong signature of the Tp<u>C</u>pN→G and Tp<u>C</u>pN→T mutations (Signatures 2 and 13) is observed in UCEC but not in COAD samples; otherwise, the MSS spectra of the two cancer types look very similar. Signatures 2 and 13 are mutational signatures of the APOBEC activity ^13^. To eliminate these differences in MSS spectra, we excluded samples with these signatures and high APOBEC enrichment values (Supplementary Table 2) from analysis (Fig. 1a, c).

Using the approach that determines the replication fork direction from data on replication timing^15–17^, we compared the rates of complementary mutations between the leading and the lagging strands for germline mutations inferred from human-chimpanzee comparison, and for somatic mutations in MSS cancers and MSI cancers (Fig. 2). In line with previous results ^16^, we found replication-driven asymmetry in MSI cancers. In general, we observed that the ratio of the rates of complementary mutations increased with the propensity of a DNA strand segment to be replicated as leading or lagging, indicating that mutation rates are asymmetric between the leading and the lagging strands. Separate analyses of UCEC and COAD cancer types provide concordant results (Supplementary Table 3), and small quantitative differences likely reflect admixture of MMR-independent mutational processes in these cancers (Supplementary Table 1). The asymmetry is very similar if we analyze only intergenic regions (Supplementary Fig. 1), suggesting that these results are not associated with transcription. We studied this asymmetry in the most extreme replication fork direction bin, because it corresponds to the genomic regions where we could predict fork polarity with the highest confidence ^17^.

**Figure 2.**
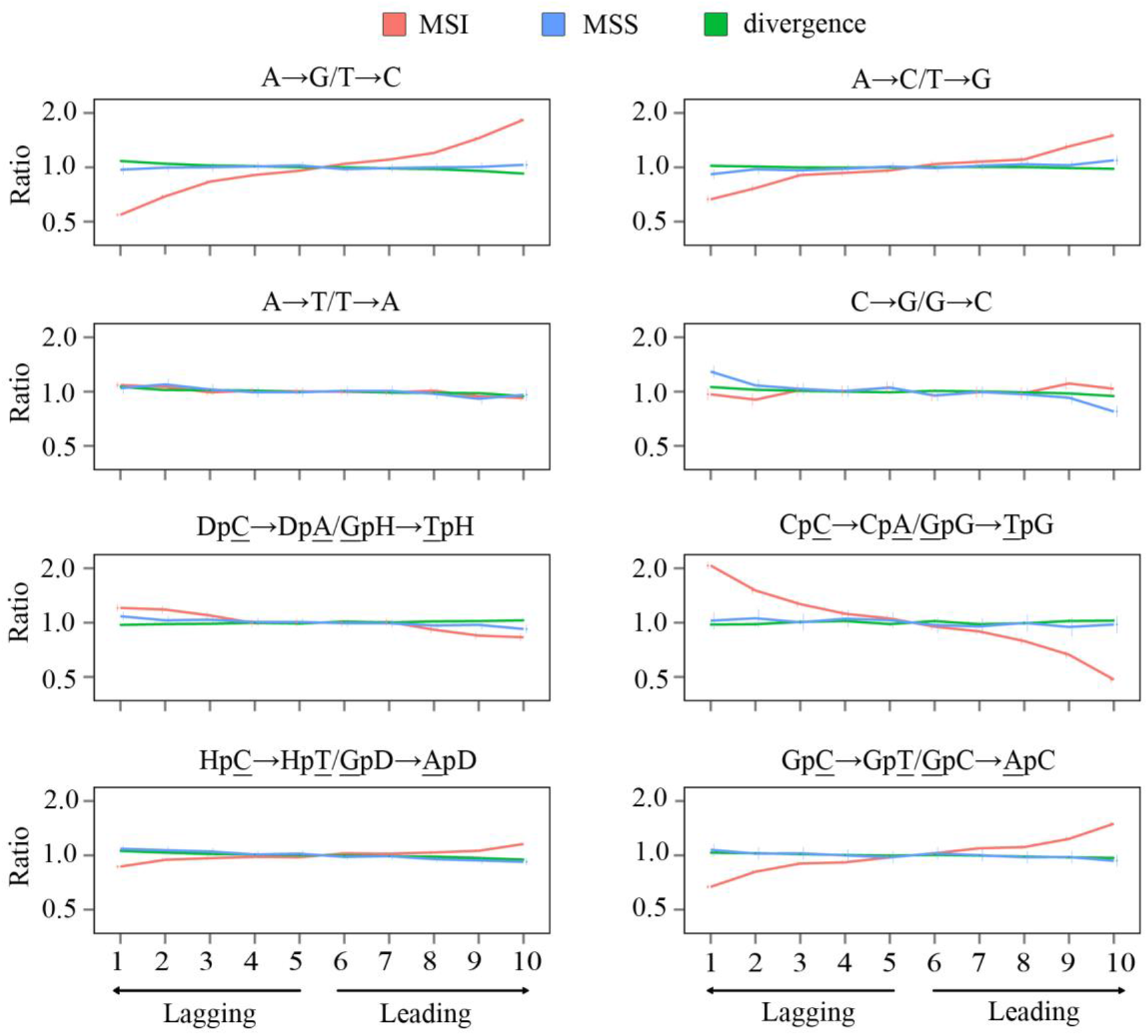
**Strand asymmetry of mutations**. Panels show ratios of the mutation rates for complementary mutations in 11 MSI (3 COAD and 7 UCEC) and 22 MSS (8 COAD and 14 UCEC) cancers, and in human-chimpanzee divergence. Horizontal axis, propensity of the replication fork to replicate the strand as lagging or leading. Vertical axis, the ratio of the frequencies of the two complementary mutation types on the strand in this category. Vertical bars represent 95% confidence intervals. D corresponds to nucleotides A, G or T. H corresponds to nucleotides A, C or T. Note the logarithmic vertical axis.

In human-chimpanzee divergence, complementary mutations are distributed nearly symmetrically between the two replicating strands, in line with previous results ^18^. Lack of asymmetry in divergence data and in MSS cancers reflects a near lack of mutation biases between the two strands in MMR-proficient cells. By contrast, in MSI cancers where the MMR system is not proficient, we observe a strongly biased distribution of complementary mutations between the leading and the lagging DNA strands. A particularly strong asymmetry (1.5-1.8-fold) is observed for mutations effectively repaired by the MMR (A→G, Cp<u>C</u>pN→A and Gp<u>C</u>pN→T). The asymmetry is small or nearly absent for mutation types depleted in mutational spectra of MSI cancers (A→T, C→G, Dp<u>C</u>pN→A, Hp<u>C</u>pN→T mutations), to the exception of the A→C mutation, where the asymmetry is 1.5-fold. In MMR-proficient cells, the observed biases have to be compensated by the MMR, which has to correct errors on one of the two strands more effectively, thus equalizing the mutation rate between strands. Moreover, to compensate for the substitution biases, the MMR has to introduce its own repair biases.

To further validate this, we reproduced our results using exome sequences of MSI and MSS cancers from the TCGA (Supplementary Fig. 2). We also analyzed exomes and genomes separately by cancer type: COAD, stomach adenocarcinoma (STAD) and UCEC. The biases observed in this bigger set of cancer types were similar and concordant between cancer types, implying that they are determined not by the cancer type but by the (in)activity of the MMR system (Supplementary Table 3).

MMR corrects single-nucleotide insertions and deletions (indels) with the highest effectiveness among all single nucleotide errors. In MMR-deficient cells, indels in homopolymer tracts and di-nucleotide tandem repeats are the most frequent type of mutations. As in yeast ^19^, in humans deletions are more common than insertions in MMR deficient cells, and their rate increases with the homopolymer tract length. We studied the most frequent deletions type, deletions of A or T in corresponding homopolymer tracts. Similarly to single-nucleotide substitutions, the efficiency of MMR in correction of deletions in homopolymer tracts may differ between the two DNA strands. We observe a small asymmetry for deletions (Supplementary Fig. 3), supporting the conjecture that the MMR activity differs between the two strands. This asymmetry has the same direction in the two cancer types, but is slightly higher in the COAD than in the UCEC.

The asymmetry in mutation rates between the leading and the lagging DNA strands is probably due to replication of these strands by different polymerases. As MMR is primarily a co-replicative process ^20,21^, we hypothesized that the observed asymmetry in MMR-deficient cancers is due to differences in types and rates of errors produced by replicative polymerases on the leading and on the lagging strands. To investigate the propensity of polymerases to particular mismatch types in humans, we employed data from patients with mutations in the exonuclease domain of one of the two major replicative polymerases, polymerase ε (polε-exo^-^) or polymerase δ (polδ-exo^-^), because the fidelity of the damaged polymerase is decreased by a factor of 100-1000, and most mutations are produced by it *^2,22^*. In particular, such mutations are frequent in cancers with inherited biallelic mismatch repair deficiency (bMMRD). We observe that the relative frequencies of different nucleotide mismatch types are radically distorted in these cancers (Fig. 3a, b). Assuming that disruption of the exonuclease domain does not affect the direction of the strand asymmetry of mutations, we can interpret these differences as mutational signatures of the corresponding replicative polymerases. We find that each polymerase usually produces one of the two complementary mutations with a higher frequency (Fig. 3 c). In particular, the biases associated with the C→A and T→G mutations are opposite: polymerase ε preferentially produces mismatches resulting in these mutations on the leading strand, while polymerase δ produces them on the lagging strand (Fig. 3c), in line with the observations in yeast *^23–25^*. Whole genomes and whole exomes data for polε-exo^-^ and polδ-exo^-^ UCEC and COAD cancers and bMMRD glioblastoma cancers demonstrate similar asymmetry patterns (Fig. 3, Supplementary Fig. 4, Supplementary Fig. 5, and Supplementary Table 4), which shows that this asymmetry is specific to the mutational processes rather than the cancer type.

**Figure 3.**
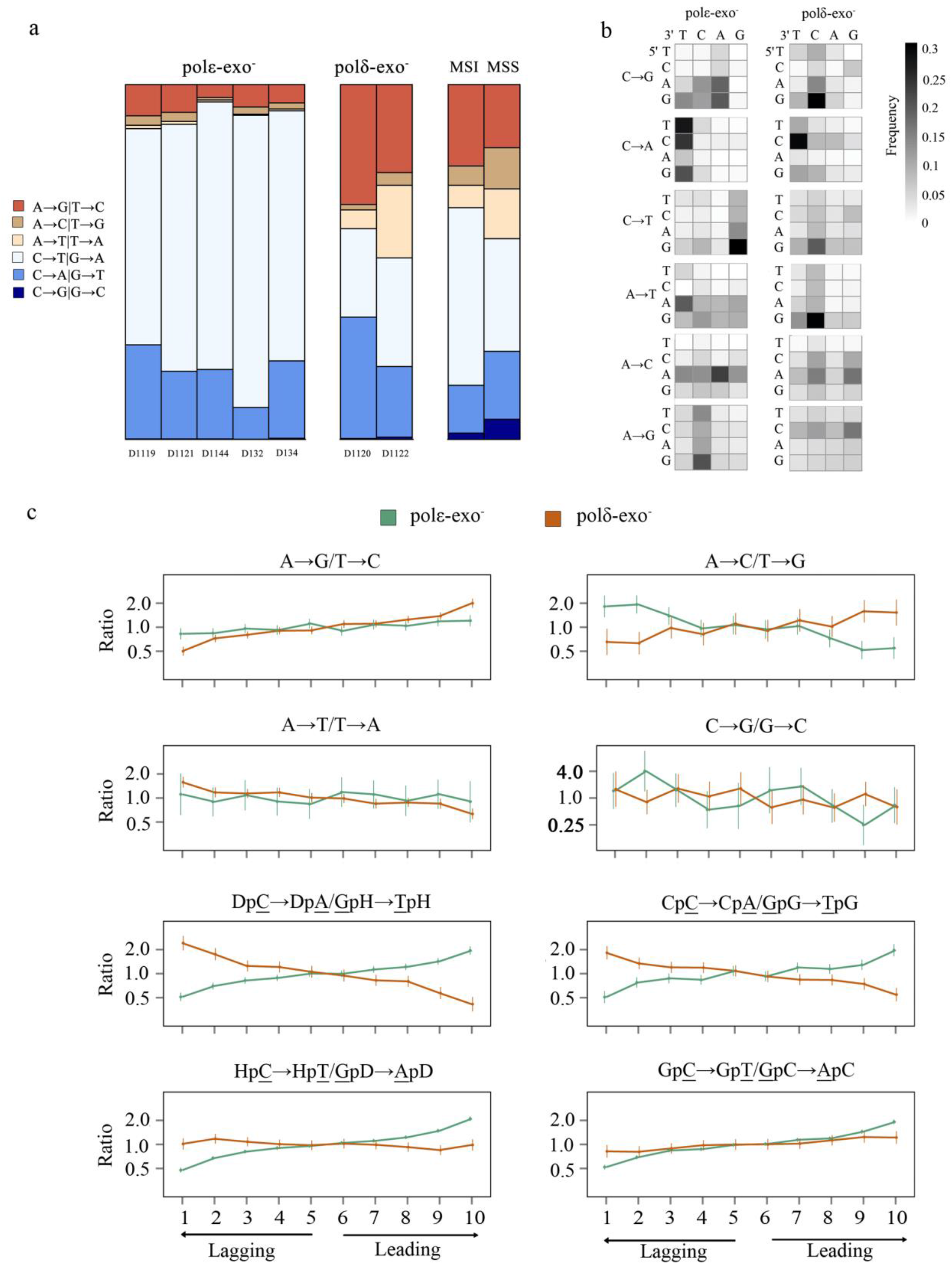
**Mutation patterns in bMMRD cancers with subsequent mutations in polε or polδ**. Data for 7 exomes of bMMRD cancer (5 polε-exo^-^ and 2 polδ-exo). a, Relative frequencies of single-nucleotide substitutions irrespective of the strand; **b**, Context-dependent mismatch frequency for each mutation type averaged between patients in each group. Vertical axis, 5’ context of the substitution; horizontal axis, 3’ context. For each single-nucleotide mutation type, the color of each of the 16 cells indicates how frequently this mutation occurs in this context; **c**, The ratio of the mutation rates for complementary mutations as a function of the propensity of the replication fork to replicate the reference strand as lagging or leading. Axes and notations same as in Fig. 2.

Notably, the direction of the mutational strand asymmetry in polδ-exo^-^ cancers generally matches that in MSI cancers with wild type polymerases (Fig. 4). We designed a model using the asymmetry observed in polε-exo^-^ and polδ-exo^-^ cancers to estimate the contributions of polε and polδ to mutagenesis in MSI cancers (see Methods). This model assumes that (i) all mutations in polε-exo^-^ (polδ-exo^-^) cancers arise from the action of polε (polδ) on the leading (respectively, lagging) strand; (ii) each of these polymerases independently contributes some fraction of mutations; and (iii) the direction of the replication fork in the 20% of the genome where replication asymmetry could be determined with the highest confidence is known exactly. We applied this model to mutation types presented in MSI spectra at high frequency (A→G, C→T, C→A). We found that the contribution of polδ to the asymmetry observed in MSI cancers is 1.9 to 4.3 fold higher than the contribution of polε (Supplementary Table 5). Accounting for the frequencies of the corresponding mutations, the overall contribution of polδ to mutagenesis was ~3-fold greater than that of polε (Fig. 4b). An independent dataset of 12 MSI COAD cancers ^26^ yielded a 2-fold greater contribution of polδ (data not shown). This implies that the main driver of the asymmetry in MSI cancers are the mutations introduced by polδ during the replication of the lagging strand.

**Figure 4.**
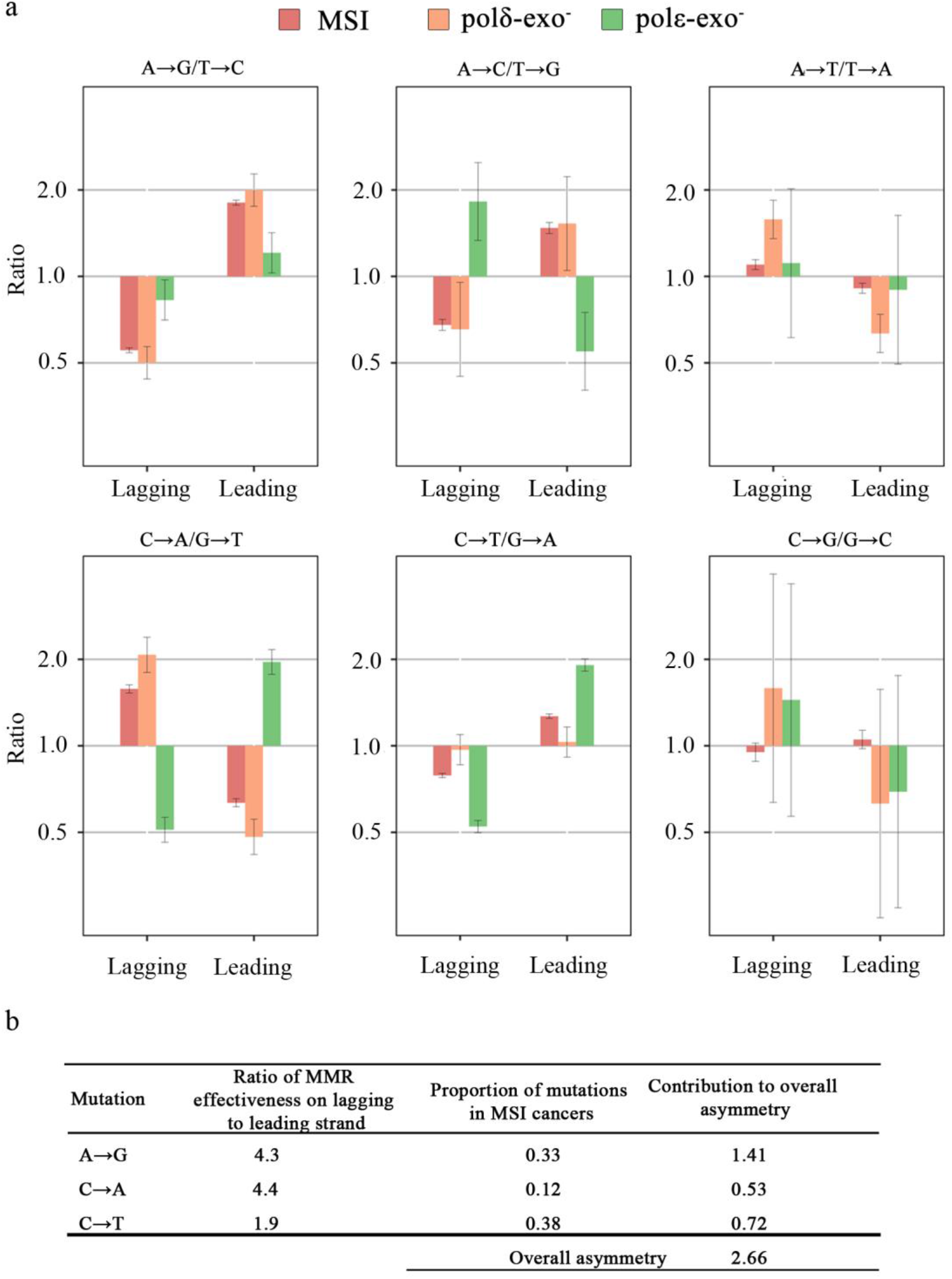
**Strand asymmetry of mutations in MSI cancers and in bMMRD polε-exo and polδ-exo cancers. a**, The asymmetry for complementary mutations in the 20% of the genome where replication asymmetry could be determined with the highest confidence (corresponding to bins 1 and 10 in Fig. 2). Error bars represent 95% confidence intervals. **b**, Model-estimated ratio of MMR effectiveness on the lagging and the leading strands.

Our analysis is subject to several caveats. Firstly, we assumed that the mutational spectra of polymerases with disrupted exonuclease domain match those of wild type polymerases. This is likely to be true: the magnitude of the strand bias is huge in yeast ^23–25^ and seems to be large in humans ^8^, and is unlikely to be radically affected by the presence of exonuclease activity. Secondly, our approach allows predicting only the preferential direction of the replication fork; therefore, the magnitude of asymmetry may be underestimated, especially if it is very strong ^17^. These caveats, however, cannot affect our conclusions qualitatively.

As no strand asymmetry is observed in wild type cells, the strand biases introduced by polymerases have to be compensated by the MMR. Since the mutations are biased towards the lagging strand, this implies that MMR is proportionally more effective in repairing mutations on the lagging strand.

Mutational spectra differed between the MMR-efficient and MMR-deficient cancers with somatic mutations in polε-exo or polδ-exo (Supplementary Fig. 6). However, the level of replication asymmetry was similar among all polε-exo^-^ or polδ-exo^-^ cancers (Supplementary Table 4). Therefore, while the MMR may change the mutational spectra, it is insufficient to compensate for the radical asymmetries introduced by polε-exo^-^ and polδ-exo^-^.

In summary, we show that the polymerase error rate is higher during lagging strand replication, that this asymmetry is primarily due to mutations produced by polδ on the lagging strand, and that these mismatches are removed more effectively by the MMR on the lagging strand (Fig. 5). This is in agreement with the biochemical property of the MMR to preferentially eliminate mismatched nucleotides on the DNA strand containing the nick ^27^. As the lagging strand is replicated in Okazaki fragments, the MMR has more time to discriminate between the template and nascent strands during its replication. More generally, the remarkable concordance between the biases in introduction and repair of mismatches helps reduce the genomic mutation rate.

**Figure 5.**
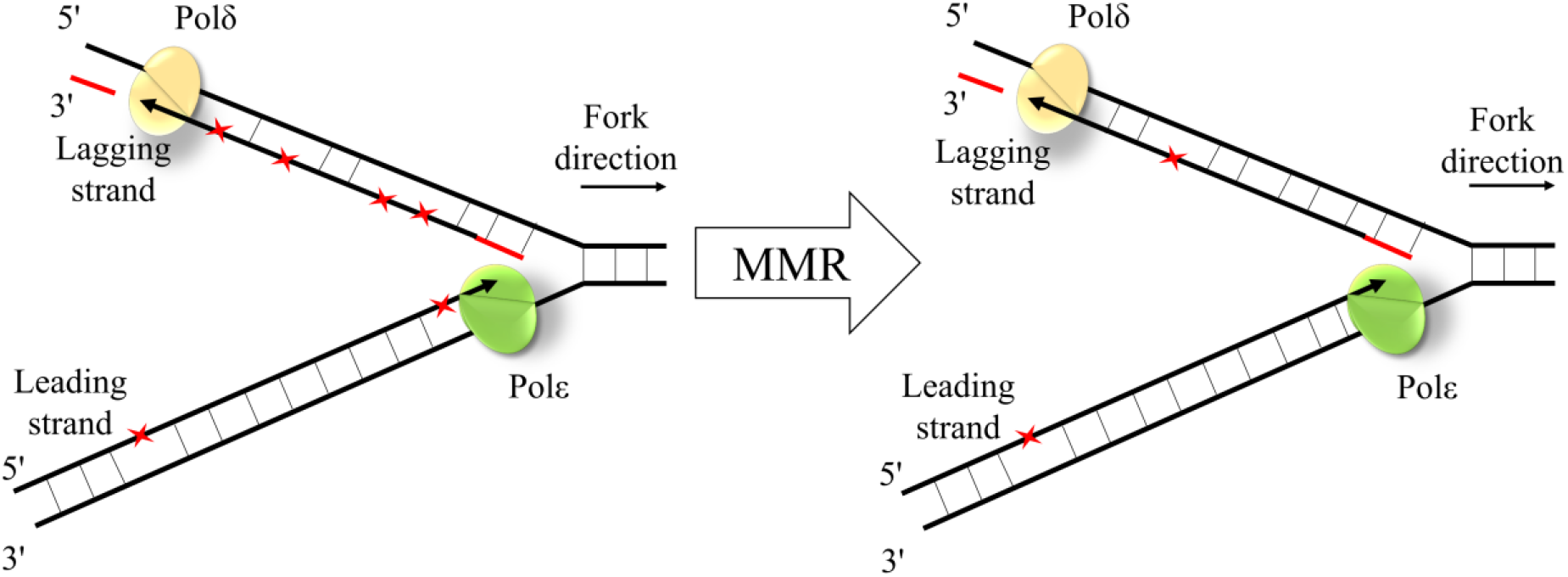
**The schematic representation of MMR effectiveness during the leading and the lagging strand replication**. While mismatches (red asterisks) are introduced more frequently during replication of the lagging strand by polδ, the lagging strand mismatches are corrected more effectively by the MMR.

## Methods

### Mutations Data

We used the following previously published data: (a) somatic mutations for whole-exome sequences of MSI (n = 159) and MSS (n = 782) cancers and for whole genome sequences of MSI (n=11) and MSS (n=27) cancers from the data portal of The Cancer Genome Atlas (TCGA) (https://tcga-data.nci.nih.gov/tcga/dataAccessMatrix.htm); (b) bam files with aligned reads for ultra-hypermutated cancers ^28^; (c) human-chimpanzee-orangutan multiple alignment from the UCSC genome browser (https://genome.ucsc.edu/). Substitution rates were calculated as the number of substitutions of a particular type divided by the number of target sites. For asymmetry analyses of the TCGA data, cancers with mutations in replicative polymerases were excluded. For analysis of the interspecies data, we obtained mutations in the human line after its divergence from the chimpanzee by maximum parsimony, using orangutan as the outgroup.

### Identification of somatic mutations in ultra-hypermutated cancers

Somatic mutations were identified using MuTect (v. 1.1.4) ^29^ under the default parameters. Mutations were then filtered against common single-nucleotide polymorphisms (SNPs) found in dbSNP and against the Catalogue of somatic mutations in cancer (Cosmic database).

### Leading vs. lagging strand asymmetry

The derivative of the replication timing at the position of the mutation was used as a proxy for the probability that the reference strand is replicated as leading or lagging in the current position, as described in ref. ^17^. The genome was categorized by these values into 10 equal bins, with the low value of the derivative corresponding to the propensity of the DNA segment to be replicated as lagging, and high value, as leading. For each bin, the numbers of substitutions and target sites were calculated. Each substitution was counted twice: as a substitution on the reference sequence with the corresponding derivative of the replication timing, and as a complementary substitution with the inverse derivative. Thus, each plot of substitutions asymmetry (Fig. 2 and 3c) is symmetric with respect to zero. Score confidence intervals were obtained for the relative risk in a 2x2 table.

### Indels analyses

We used data on single nucleotide deletions in poly-A and poly-T tracts for MSI cancer genomes, for tracts with length six to eight identical nucleotides where enough deletions were found for analysis.

### APOBEC enrichment

APOBEC enrichment was counted for each sample as ratio of C→K mutation rates in TpCpW and VpCpW contexts. Weigths of mutational signature were calculated using R-package “deconstructSigs” ^30^.

### Model for mutational biases between strands

We calculated the ratios of the mutation rates using the following logic. From Fig. 4, for each type of mutation A→B, we obtained the ratio of its rate r(A→B) and the rate of its complement r(A′→B′) on the leading strand. Then

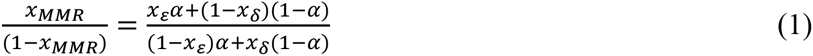

where α is the fraction of mutations A→B and A′→B′ that are produced by polε on the leading strand; 1-α is the fraction of such mutations that are produced by polδ on the lagging strand; xε and 1-*x*_ε_ are the fractions of mutations A→B and A′→B′ respectively produced by polε; *x*δ and 1-*x* δ are the fractions of mutations A→B and A′→B′ respectively produced by polδ; and *x*_MMR_ and 1-*x*_MMR_ are the fractions of mutations A→B and A′→B′ respectively on the leading strand in MSI cancers.

For example, consider mutation C→A/G→T. From Fig. 4, *x*_MMR_/(1-*x*_MMR_) is 0.65. As the C→A/G→T ratio for the leading strand in polε-exo^-^ cancers is 1.96 (Fig. 4), the fraction of C→A mutations produced by polε is *x*_ε_= 0.66, and the fraction of G→T mutations produced by polε is 1-*x*_ε_= 0.34. Similarly, as the C→A/G→T ratio for the lagging strand in polδ-exo^-^ cancers is 2.07, the fraction of C→A mutations produced by polδ is *x*δ= 0.67, and the fraction of G→T mutations produced by polδ is 1-*x*_δ_= 0.33. From equation 1, (1-α)/α = 4.0. In other words, for this mutation type, polδ produces mismatches leading to this mutation type on the lagging strand ~4 times more often than polε produces them on the leading strand. As no strand bias is observed in MSS cancers, the repair bias by MMR has to be exactly inverse, repairing polδ mutations 4 times as effectively as polε mutations. The results for all 5 mutation types are calculated similarly (Table S3). The overall strand asymmetry of MMR was calculated as the mean asymmetry across the five mutation types, weighted by the proportions of each mutation in MSI cancers.

## Acknowledgements

We thank Shamil Sunyaev, Pasha Mazin and Sonya Garushyants for useful discussion. This work was performed at IITP RAS and supported by the Russian Science Foundation grant no. 14-50-00150.

## Author Contributions

V.B.S. designed the study, M.A.A. performed analyses, M.A.A., G.A.B. and V.S.B. wrote the paper, S.I.N. provided data for cancer samples and assisted in interpretation of results and writing the paper.

The authors declare no competing financial interests.

